# The spore of the beans: Spatially explicit models predict coffee rust spread in fragmented landscapes

**DOI:** 10.1101/2020.10.16.343194

**Authors:** E.M. Beasley, N. Aristizábal, E. Bueno, E.R. White

## Abstract

Landscape structure influences the spread of plant pathogens, primarily by affecting pathogen dispersal. Coffee leaf rust (*Hemileia vastatrix*), a fungal disease that causes heavy economic losses in the coffee industry, is likely to be affected by landscape structure via dispersal of its wind-borne spores. Previous studies have found positive associations between leaf rust incidence and the proportion of pasture cover, suggesting that deforestation may facilitate rust spore dispersal. We explored this idea by modeling the spread of rust transmission in simulated landscapes. Specifically, we modeled within-patch transmission using a probabilistic cellular automata model, and between-patch transmission using a random walk with spore movement inhibited by forest canopy cover. We used this model to understand how the spread of coffee rust is affected by: 1) clustering of coffee plants, 2) clustering of deforestation, and 3) proportion of landscape deforestation. We found that clustering of coffee plants is the primary driver of rust transmission, affecting the likelihood and severity of rust outbreak. Deforestation is important in landscapes with high clustering of coffee: rust outbreaks are more severe in landscapes with a higher proportion of deforested areas, and more variable in landscapes where deforested areas are more evenly dispersed throughout the landscape.

## Introduction

Many ecological systems are characterized by their landscape configuration, including natural or anthropogenic habitat fragmentation (Levin 1992; Fahrig 2003). Landscape structure includes the distribution and quality of habitat, which in turn affects the connectivity of habitat patches. Furthermore, landscape structure is known to affect various species through its effect on connectivity of high quality habitat patches, patch size and extinction risk, as well as edge effects (Wiegand et al. 2005; Gavish et al. 2012; Fahrig 2013; Haddad et al. 2015; White and Smith 2018). One area where landscape structure has been particularly relevant is in the spread of disease (White et al. 2018).

The conversion of native habitat for agriculture and urban development is associated with an increase in infectious diseases (Ellwanger et al. 2020, Gibb et al. 2020). Although evidence remains mixed (Hagenaars et al. 2004; Tracey et al. 2014), a lot of work suggests that both habitat fragmentation and decreased habitat quality can increase the likelihood of disease spread (White et al. 2018). Plantagenest et al. (2007) suggest there are four landscape-level factors that influence plant pathogen dynamics: 1) landscape composition influences global inoculation pressure, 2) landscape heterogeneity impacts pathogen dynamics 3) landscape structure affects pathogen dispersal, and 4) landscape properties can induce the emergence of pathogens. Yet, there have been limited efforts to understand mechanistically how landscape structure affects plant populations (Cunniffe et al. 2015a).

Here we use the plant disease, coffee leaf rust (*Hemileia vastatrix*), as a model system for understanding issues of fragmentation and disease spread. First recorded in 1879, coffee leaf rust is a fungal disease notably recognized for completely destroying the coffee industry in Sri Lanka, which used to be one of the largest coffee-producer countries in the world. Since the 1970s, coffee rust has spread to the largest coffee-producing regions in the world including Brazil, Mexico, and Colombia. Reports of up to 30-50% losses due to coffee rust in Brazil and Costa Rica, 31% in Colombia, and 16% in Central America make this disease an urgent priority for the coffee-growing industry (Baker 2014; Avelino et al. 2015; McCook and Vandermeer 2015; Zambolim 2016; Cerda et al. 2017).

*Hemileia vastatrix* is an obligate fungal pathogen affecting cultivated coffee species including *Coffea arabica* and *Coffea canephora. H. vastatrix* primarily infects leaf tissues leading to defoliation and reductions in vegetative growth all of which reduce coffee berry yields (Waller 1982). Infections are commonly recognized by the formation of orange pustules (uredina) underneath leaves that release between 300,000 to 400,000 spores into the environment (Kushalappa and Eskes 1989). The infection process is composed of three principal steps: germination, penetration, and colonization of spores, with germination and penetration requiring the presence of running water (Kushalappa and Eskes 1989). Other factors influencing germination and penetration include leaf age and spore concentration and distribution at the infection site. Colonization of spores into the stomatal opening of the leaves is followed by the production of spores (sporulation). Within one to three weeks, pale yellow spots on the underside of leaves are the first signs of infection. Depending on the age of the leaves, spore production can begin anywhere from two weeks to months following infection (Kushalappa and Eskes 1989). At maturation, spores are released into the air and can travel as high 1000 meters above coffee canopies (Martinez et al. 1975). Disease severity has been associated with the number of visible spores per leaf, the duration of leaf wetness, rainfall, and minimum temperatures (Zambolim 2016). At the canopy level, rainfall can facilitate the spread of spores via raindrops that splash between leaves (Kushalappa and Chaves 1980; Waller 1982). Thus, the spread and severity of coffee leaf rust are heavily determined by climatic conditions.

In addition to rain, spore dispersal is primarily facilitated by strong wind patterns that potentially carry spores hundreds of miles from the origin (Waller, 1982). Earlier studies have provided evidence for the dispersal of airborne spores through trapping techniques above and within coffee tree canopies (Martinez et al. 1975, Becker et al. 1975). Under outbreak conditions, maximum wind speed and low relative humidity have been associated with increases in spore dispersal during late morning and afternoon (Becker et al. 1977a). Furthermore, wind speeds between 12-20 km/hr have been linked to high numbers of spores with moderate numbers at wind speeds as low as 7 km/hr (Martinez et al. 1977). Wind gusts, in combination with rainfall, can indirectly affect the dispersal of spores by reducing the size of rain droplets that land on coffee plants under dense shade conditions (Boudrot et al. 2016). Conversely, under non-shade or open canopy conditions, wind gusts are an important driver for spore dispersal, highlighting the complexity of dispersal patterns across space (Boudrot et al. 2016).

Although there is a vast amount of knowledge on the epidemiology and environmental drivers of coffee rust, the impacts of landscape structure on coffee rust spread remain unknown. Local landscape context such as shade cover and proximity to pasture have been shown to be positively correlated with coffee rust incidence, highlighting the need to investigate how habitat fragmentation due to deforestation influences the spread of coffee rust (Avelino et al. 2012). Yet, few studies have focused on how landscape patterns may influence the spread and infection rates of coffee rust. Therefore, here we use simulation models to investigate how landscape composition and configuration influence the spread of the coffee rust.

We hypothesize that the windborne dispersal of rust spores is facilitated by landscape composition and configuration. Specifically, we examine the effects on disease transmission from the clustering of coffee plants, proportion of deforestation within the landscape, and the degree to which deforested areas are scattered in space. We predict that rust spores will disperse more readily through landscapes with high coffee clustering and deforestation levels; resulting in a higher incidence of coffee rust in landscapes that exhibit these characteristics.

## Methods

### Landscape simulation

We modeled landscapes using two 100 x 100 grids constructed using the package NLMpy (Oliphant 2006) in Python 3.7.1 (Python Software Foundation 2018). Landscapes had reflective boundaries and two landscape characteristics. The first aspect in each landscape represents the presence or absence of coffee plants and was constructed using the NLMpy function randomClusterNN (Etherington et al. 2015) which is an adaptation of the modified random cluster algorithm (Saura and Martinez-Millan 2000). We controlled aggregation of the coffee plants by a parameter ranging from 0.1-0.3, with 0.3 being the most clustered (Table 1).

**Table 1.** Parameter values for simulated landscapes.

| Parameter | Values |
| --- | --- |
| Coffee clustering | 0.1, 0.2, 0.3 |
| Proportion of deforestation | 0.15, 0.3, 0.45, 0.6, 0.75 |
| Clustering of deforestation | 1, 2, 3, 4, 5 |

The second landscape aspect represents the surrounding matrix and “mirrored” the coffee array (i.e. cells containing coffee were empty in the matrix array). Values in each matrix cell represented canopy density. The proportion of the array consisting of deforested cells, or cells with a canopy density less than 30%, ranged from 0.15-0.75. Clustering of deforested cells was controlled by a parameter taking values from 1-5, with 1 being the most clustered and 5 the most dispersed (Table 1).

We generated 50 replicate landscapes for each combination of parameter values for a total of 3,750 simulations. We initialized rust infection in one randomly selected coffee cell in each simulated landscape.

### Modeling rust transmission

We modeled transmission of coffee rust through the simulated landscapes using a two-step process linked by two transition processes (Figure 1). The two processes in our model reflect differences in rust dispersal mechanisms at the local (within-patch) and regional (between-patch) scale. Local transmission of spores can occur through wind (Kushalappa and Eskes 1989), the impact of raindrops of coffee leaves (Rayner 1961a, b; Boudrot et al. 2016), and leaf-to-leaf contact (Vandermeer et al. 2018). For all of these local mechanisms, transmission is thought to decline with increasing distance from the infected source (Vandermeer et al. 2018); therefore a coffee plant with more infected neighbors is more susceptible to infection than a plant with few or no infected neighbors. Regional transmission, by contrast, is primarily through wind (Kushalappa and Eskes 1989; Avelino et al. 2015) and dispersal at the landscape scale is more likely to be affected by regional wind patterns and barriers inhibiting wind movement.

**Figure 1.**
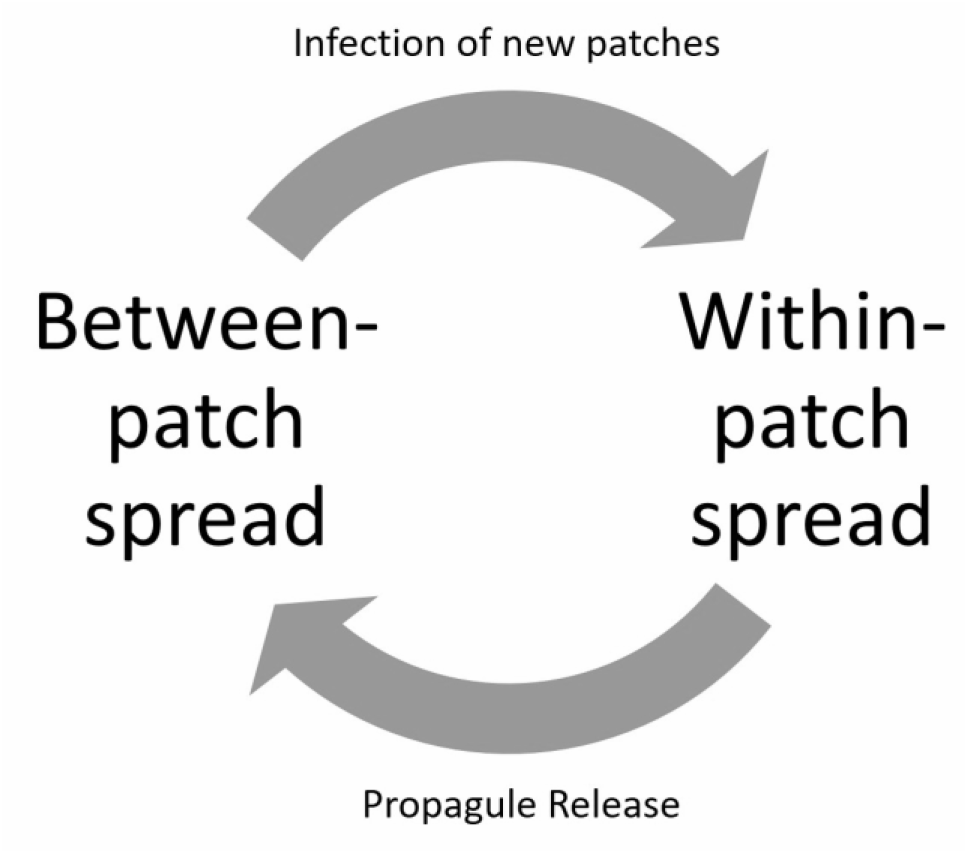
Conceptual diagram of the two-step model and transition processes.

We modeled local transmission using a stochastic cellular automata model (Wolfram 2002), in which the probability of infection in the focal cell at time *t+1* is determined by *p* ~ *Beta*(*N, 8 – N*); where *N* is the number of infected cells in the focal cell’s Moore neighborhood. Because the model assumes that local spread only occurs via transmission between immediate neighbors, a cell with no infected neighbors had a probability of infection *p* = 0.

After modeling local spread, the model transitioned to regional spread when all infected coffee plants at the edge of a patch released 1 spore into an adjacent, randomly selected landscape tile. After new spores were released, all spores moved throughout the landscape using a modified random walk. At the beginning of the walk, all spores were assigned equal “movement values” which determined how far the spore could move during the time interval. For each movement, the target landscape tile was randomly selected from the focal cell’s Moore neighborhood, and the spore moved into the target tile. We subtracted the canopy density in the target cell from the spore’s movement value. We repeated the process for all spores with a positive movement value, until the movement value for all spores was less than or equal to 0.

Upon completion of the simulated walk, each spore adjacent to an uninfected coffee plant infected the plant with a probability of success set at 0.5. If a spore was adjacent to multiple uninfected plants, the target plant was selected randomly. Spores which successfully infected a plant were removed from the simulation. We repeated this four-part process for 1000 time steps per simulated landscape.

### Analyses

We calculated landscape-level rust prevalence at the final time step of each simulation. Assuming the distribution of prevalence values followed a beta distribution, we estimated *α* and *β* using the R package fitdistrplus (Delignette-Muller and Dutang 2015; R Core Team 2020). Using the estimated values of these parameters, we calculated 1) the expected value of the distribution, 2) skewness, and 3) the concentration parameter *κ*, which describes the width of the distribution around the mode. We also compared the maximum prevalence of each distribution and performed these calculations with 1) all simulation replicates pooled, 2) replicates separated by clustering value, and 3) replicates separated by all parameter values. We evaluated associations between deforestation, dispersion, and the properties of the beta distribution using Spearman’s *ρ*. Correlations of *ρ* ≥ 0.4 and *ρ* ≤ −0.4 were considered strong associations.

## Results

Final rust prevalence across all parameter values ranged from 0.1–55.0% (Figure 2). The estimated parameters of the full distribution were estimated at *α* = 0.855 and *β* = 10.564. These values correspond to an expected value *E*(*x*) = 0.075, a skewness of 1.697, and concentration parameter *κ* = 11.419.

**Figure 2.**
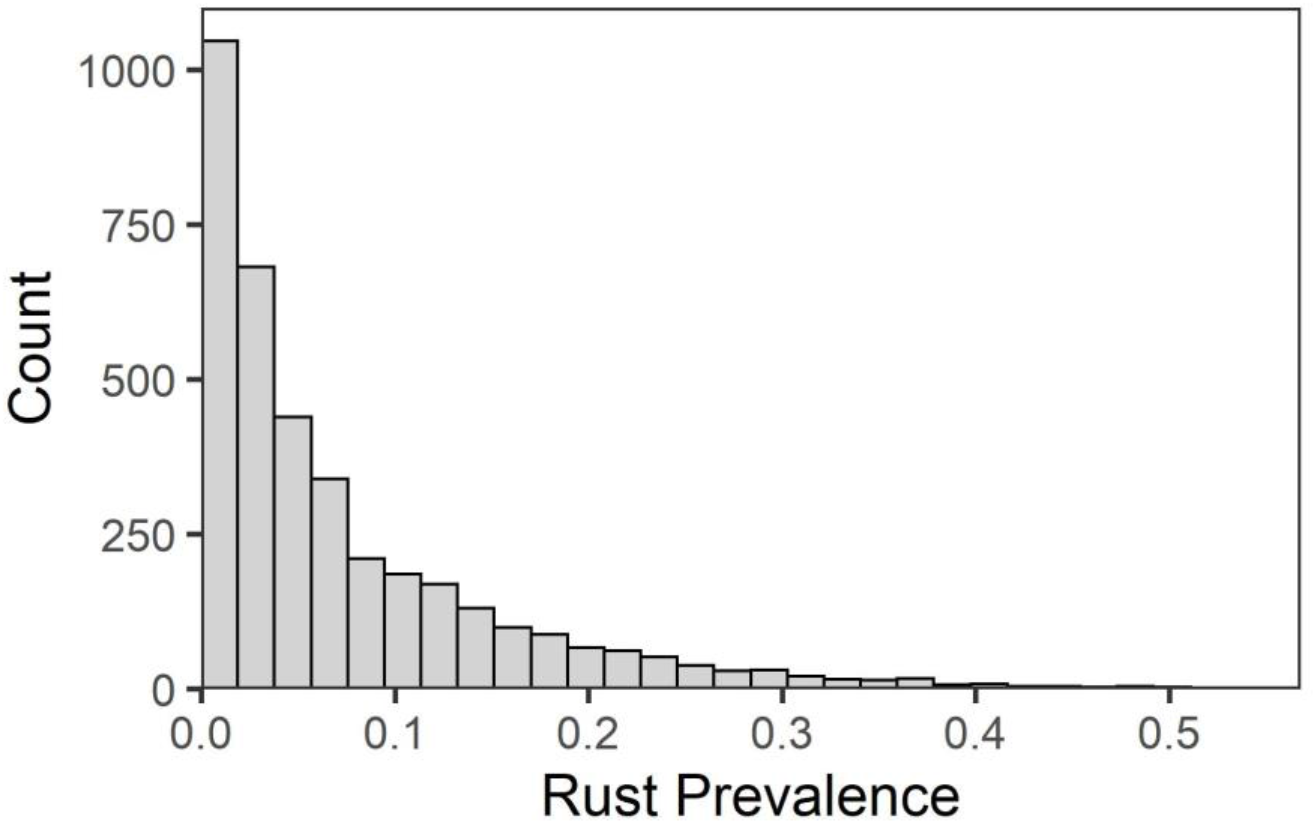
Distribution of model outcomes across all parameter combinations. Assuming the data follow a beta distribution, the expected value is 0.075, the skewness 1.697, and the concentration parameter *κ* is 11.419.

All four metrics varied among coffee clustering values. The expected value and maximum infection was greatest in landscapes with high clustering, but did not appear to differ between low and moderate clustering values (Figure 3A, 3B). Conversely, the distributions of outcomes at high clustering values were less skewed and had lower values of precision (*κ*) than distributions at low or moderate clustering values (Figure 3C, 3D).

**Figure 3.**
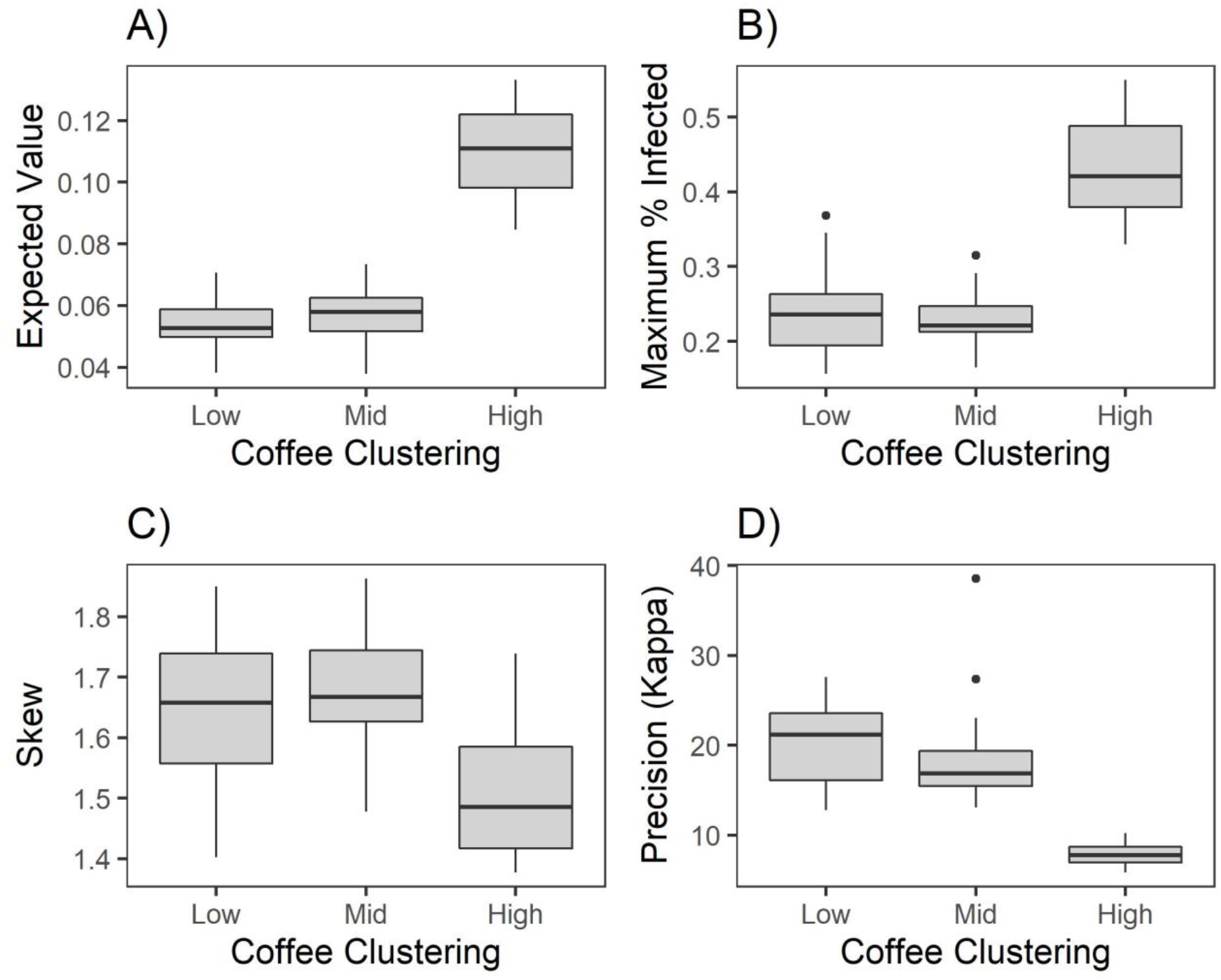
Different values of coffee clustering yield distributions of rust prevalence that vary in shape. Simulations in which coffee was highly clustered resulted in distributions with a greater expected value (i.e. typical outbreak size, panel A) and a greater maximum value (B). High clustering of coffee also resulted in distributions that were less right-skewed (C) and had more variability (D).

We did not find consistent effects of deforestation and dispersion within clustering values; and the strongest patterns were typically present at the highest value of coffee clustering. The expected value of infection tends to increase with deforestation (*ρ* = 0.510, Figure 4A). Maximum infection tends to decrease with dispersion, but this association is weak (*ρ* = −0.255, Figure 4B). Deforestation was also strongly associated with skew, with a greater degree of right skew at low values of deforestation (*ρ* = −0.514, Figure 4C). Finally, the concentration parameter tended to be highest at high values of dispersion (*ρ* = 0.455, Figure 4D). The concentration parameter was also strongly associated with dispersion at moderate values of coffee clustering (*ρ* = 0.412, Figure 5), a pattern which held even when outliers were removed (*ρ* = 0.402).

**Figure 4:**
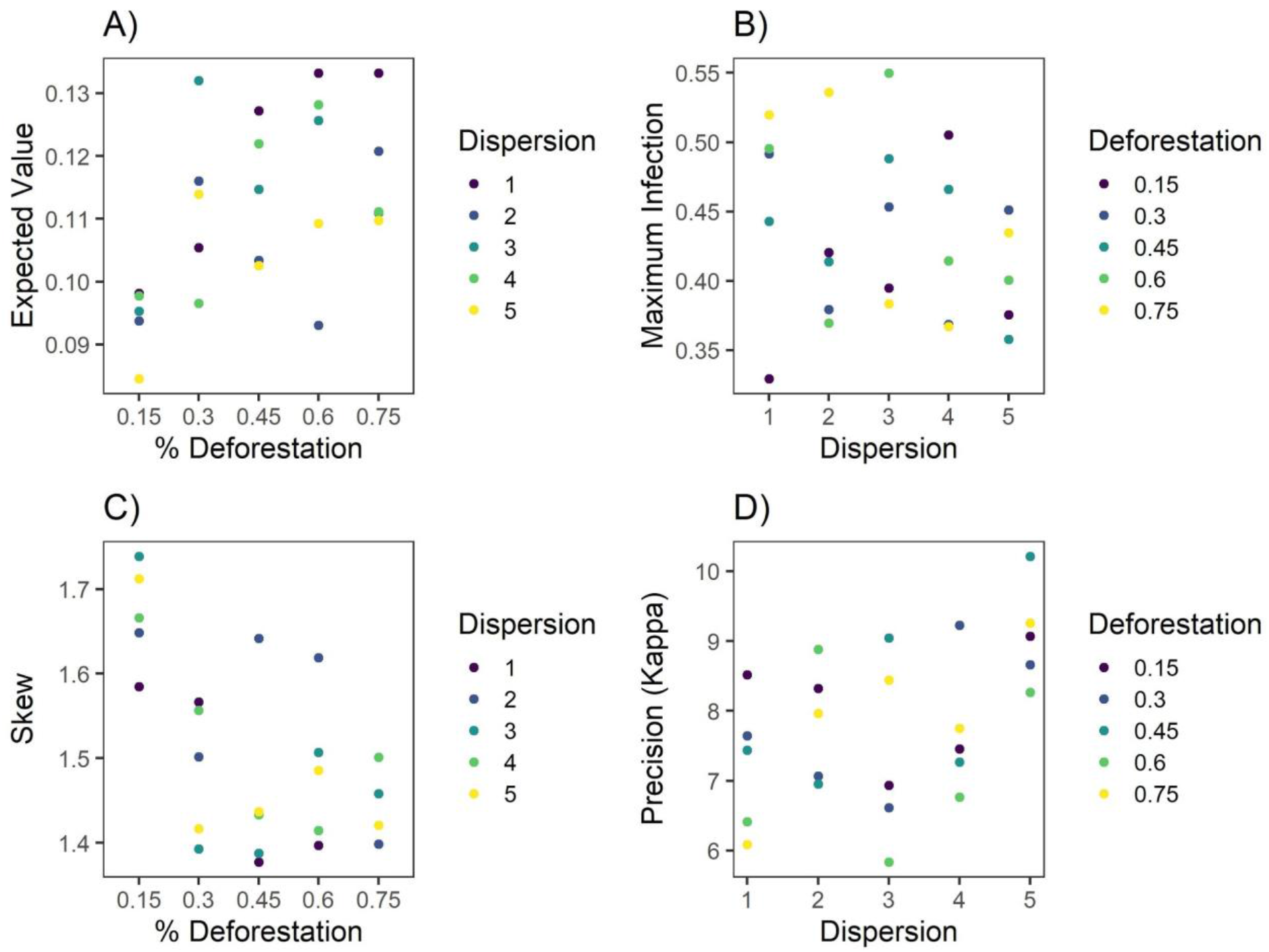
Effects of deforestation and dispersion appear at the highest value of coffee clustering. When a greater proportion of the landscape is deforested, the typical outbreak size tends to be larger (A) and the distribution less right-skewed (C). When deforested areas are dispersed more evenly throughout the landscape, maximum rust infection tends to be lower (B) and outbreak size more predictable (D).

**Figure 5:**
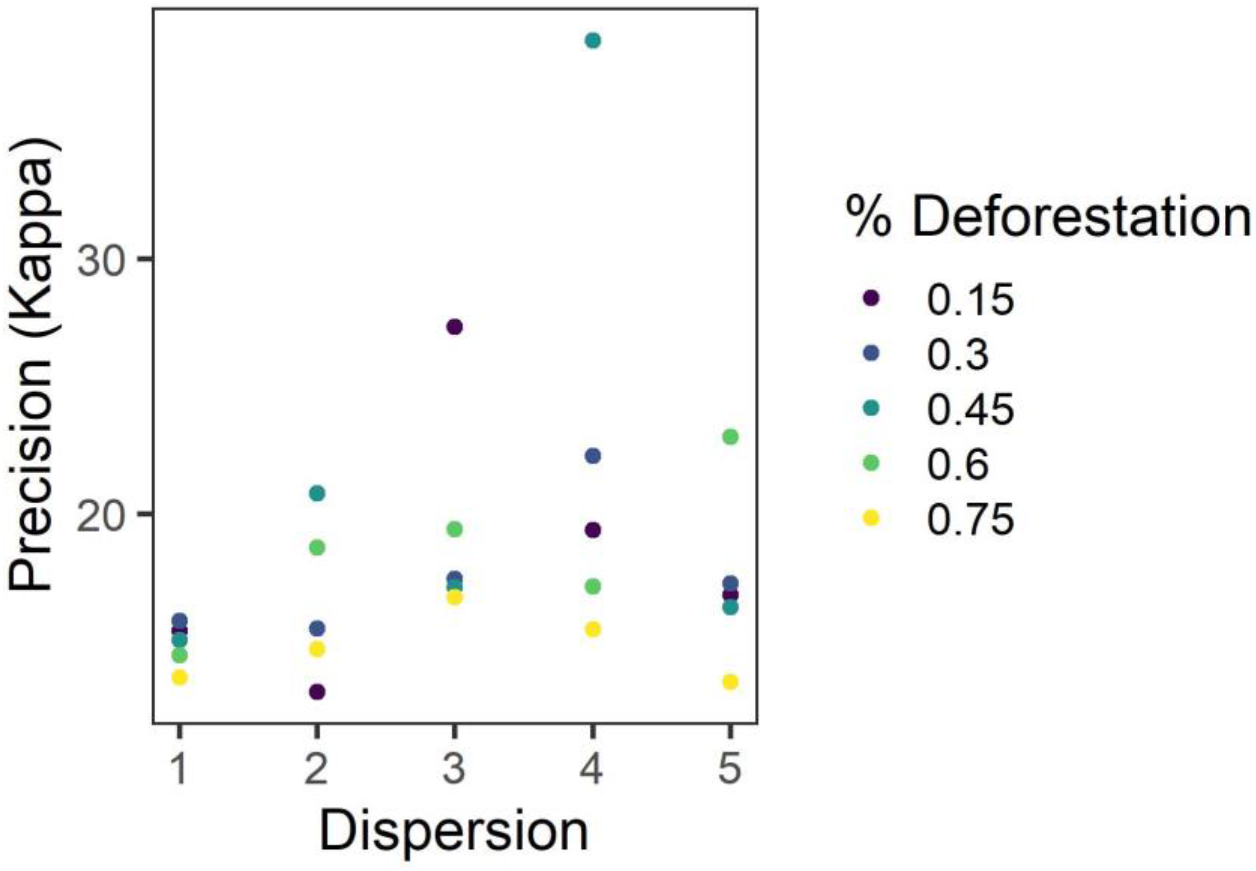
At the moderate value of coffee clustering, the spread of the distribution tends to increase with increased dispersion of deforested cells (*ρ* = 0.412). This pattern remains when outliers are removed (*ρ* = 0.402).

## Discussion

Generally, in our simulated landscapes rust outbreaks were fairly localized, with most outbreaks small in scale. Of the three landscape metrics that we manipulated (coffee clustering, deforestation, and dispersion of deforested cells), coffee clustering is the major driver of coffee rust spread (Figure 3). At the landscape level, coffee clustering leads to more coffee rust and higher variability in the outbreak size (Figure 3). The amount of forest and how the forest is distributed in the landscape become important predictors of coffee rust in landscapes with high degrees of coffee clustering. When coffee is highly clustered, coffee rust prevalence increases with deforestation (Figure 4A) and its variability decreases the more deforested areas are scattered throughout the landscape (Figure 4D).

### Deforestation and coffee rust management at the farm level

Our simulation results suggest that clustered coffee plants may act as significant sources for rust spread in deforested areas. Conventional management practices mainly depend on the foliar and systemic application of fungicides, including copper oxides, copper hydroxides, triazoles, and strobilurin (Capucho et al. 2013; Zambolim 2016). The timing of chemical controls are typically determined by weather, calendar day, plant age or via disease monitoring-based methods (Capucho et al. 2013). The development of coffee rust resistant varieties also provides farmers with protection against outbreaks (Avelino et al. 2004). However, reductions in resistance durability has arisen due to the appearance of highly virulent rust strains (Silva et al. 2006).

Crop management practices such as pruning, fertilizing, manipulation of crop density and the addition of shade trees can have complex effects on spore dispersal and germination (Avelino et al. 2004, 2006). For instance, when rainfall levels are low, shade trees protect coffee plants by reducing the amount of raindrops falling onto leaves, thus reducing the dispersal of spores (Jaramillo and Chaves 1998). On the other hand, when rainfall levels are high, shade trees can have opposite effects on dispersal by acting as “living gutters” that create large droplets that fall onto coffee trees and stimulate spore dispersal. Shading practices can also reduce wind speed, resulting in reductions in aerial spore dispersal (Jaramillo and Gomez 1989). More recently, Gagliardi et al. (2020) demonstrated the importance of shade tree leaf and canopy traits on wind speed dynamics and spore dispersal at the edges of coffee agroforestry systems. Shade tree traits such as leaf thickness, canopy base height, and canopy openness were found to be important for reducing throughflow wind speeds and for promoting spore settling at the interior edge of coffee plots. High density planting of coffee plants, which promotes self-shading, protects coffee trees in the same manner as shade trees, but can be detrimental as increases in leaf area index can facilitate spore establishment (Arcila and Chaves 1995). These studies highlight the complex interactions between coffee management practices and microclimatic factors in driving or reducing the risk of coffee rust outbreaks at the farm level. To this end, the outcomes from our simulations may have direct implications for rust management practices as high clustering of coffee plants resulted in larger and less predictable outbreaks.

Understanding the socio-ecological factors involved in local spatial configuration of crop plants is crucial to reduce susceptibility to plant disease dynamics. Hajian-Forooshani and Vandermeer (2020) compared simulated coffee rust spread at the farm-level following a simple null model to a network model from empirical data to understand the spatial structure of coffee rust dynamics. Complementary to our results, they found that shade reduction and uniform coffee plot arrangements likely increases the probability of coffee rust outbreaks. Thus, spatial arrangement of coffee plants is highly important to consider either when a new coffee crop is being established or when an individual coffee plant is replaced.

As coffee plants age, they decrease their production and get replaced. When this opportunity to restructure a coffee plot arises, our study suggests that in addition to avoiding uniformity (Hajian-Forooshani and Vandermeer 2020), coffee plots should be less clustered to decrease the spread of coffee rust. This is in line with disease modeling of other systems, where clustering of individuals leads to more frequent and severe outbreaks (Lieu et al. 2015; Althouse et al. 2020).

One method to prevent coffee clustering is incorporating shade crops, an already commonly used strategy to manage coffee rust. Adding Musaceae species (banana and plantain) or *Erythrina poeppigiana* (also known as poró) to break the wind reduces spores dispersal, but it can also help reduce the clustering of coffee plants. Shade trees exhibit great variability and *porós* are regularly pruned as part of the management, but the most effective shade trees are dense porós with thick leaves. However, this reduction of wind speed comes with a trade-off as coffee rust spores settle more at the edges of a coffee plot (Gagliardi et al. 2020). Combined plot-level strategies can be beneficial and the investment of replacement of individual plants with coffee rust-resistant varieties placed at the edge of a coffee plot.

### Deforestation and coffee rust management at the landscape level

The emergence of coffee rust in Brazil during the 1970’s had a large impact on landscape management across Latin America, as farmers recognized the importance of shade trees as a means to control coffee rust outbreaks. The industrialization of coffee in combination with the appearance of coffee rust has transformed the landscape of many coffee growing regions, creating a “patchy” landscape composed of conventional and traditional “shade farms” (Rice 1999). Moreover, the homogenization of coffee landscapes, primarily through the clearing of forested areas, has been previously linked to coffee rust outbreaks and continues present major challenges for disease control (Avelino et al. 2004, 2006; Boudrot et al. 2016; Perfecto et al. 2019). To this end, deforestation, and thus the loss of shade tree protection, can facilitate the dispersal of rust spores by allowing wind gusts to infiltrate coffee canopies (Perfecto et al. 2019). Our results agree with these mechanistic explanations for the spread of coffee rust within a simplified-homogenous landscape as high levels of clustering and deforestation resulted in higher prevalence of coffee rust infections. Thus, efforts to undo the effects of landscape homogenization through conservation and reforestation practices may serve as an effective approach to manage coffee rust at the landscape level.

Localized management strategies are insufficient for managing many fungal diseases due to the dispersal of windborne spores. Previous farm-scale management tactics, such as host removal through culling or planting resistant varieties, fail to contain epidemics because they underestimate the spatial scale of the outbreak (Gilligan et al. 2007). It is possible that high-profile failures to contain plant disease outbreaks, such as sharka (Rimbaud et al. 2015) and citrus canker (Irey et al. 2006) are due to implementing local, reactive strategies rather than a landscape-level approach (Gilligan et al. 2007; Fabre et al. 2019). Our study suggests that localized strategies are particularly ineffective in highly connected landscapes (Figure 3), where coffee patches are large and clustered together. This finding reflects previous studies regarding disease transmission and landscape modality (Macfadyen et al. 2011). Characteristics of the matrix surrounding farmland, specifically deforestation, can also facilitate or inhibit disease spread (Figure. 4). This demonstrates the need for management strategies that address processes occurring at multiple scales (Amico et al. 2020).

The majority of the coffee in the world is produced in small farms of less than 10 hectares (Jha et al. 2014). Therefore, cooperation between farmers is critical for managing coffee rust outbreaks at the landscape level. For instance, communication among neighboring farmers may help minimize clustering of coffee plants and facilitate community-based reforestation efforts. Additionally, the exchange of disease monitoring and management information may help contain the spread of coffee rust at the landscape level. Examples of cooperation among farmers have been documented for the management of plant diseases such as cassava brown streak disease and Huanglongbing (Bassanezi et al. 2013; Legg et al. 2017). Coordinated management is thought to be most successful among farmers and their immediate neighbors, but at larger spatial scales, competing interests and values may hinder collective management strategies (Sherman et al. 2019). An integrative approach that incorporates on-farm, neighborhood, and landscape management strategies may serve as a productive way for farmers, land managers, and government officials to collectively manage coffee rust (Amico et al. 2020).

### Mathematical modeling and simulation of landscape processes

The relative simplicity of our model allows us to focus on landscape effects without having to filter through “noise” created by other environmental processes. However, this simplicity comes with the disadvantage of omitting a handful of factors that may influence how well our model reflects reality. One such factor is the finite landscape boundaries present in our simulated landscapes. Finite boundaries are frequently a necessity in simulated landscapes due to limited computation time, but tend to bias movement-based simulations (Koen et al. 2010). This problem is most evident in cells adjacent to the landscape boundary, where moving agents have a more limited selection of cells they can potentially enter (Keane et al. 2006). While our simulation does not include measures to mitigate boundary effects, such as a buffer around the landscape (Koen et al. 2010), a visual inspection of the outcomes of our model suggests that simulations where rust infection starts near the edge do not yield unusually high or low values of rust prevalence.

Our model does not include the effects of wind, sunlight, and humidity, all of which are known to affect rust transmission and germination (Kushalappa and Eskes 1989; Avelino et al. 2015).

Wind, in particular, is thought to be the primary mechanism driving landscape-level dispersal of rust spores. By omitting wind, it is likely our model 1) underestimates the importance of deforestation as a driver of landscape-level prevalence, and 2) fails to detect effects of starting conditions (i.e. location of initial infection) on final prevalence. Despite these faults, our model successfully outlines broad patterns of rust spread and the landscape factors influencing outbreaks.

Modeling plant disease spread presents unique challenges (Cunniffe et al. 2015a). For instance, in contrast to human diseases, plants are sessile and data on the location and the disease state of individuals can be particularly difficult to obtain (Cunniffe et al. 2015a). However, simulations and mathematical modeling can play a major role in plant disease management because most issues regarding management are spatial in nature (Fabre et al. 2019). This is particularly true for pathogens that can disperse long distances because processes at local and landscape scales can influence their spread (Plantegenest et al. 2007; Gilligan 2008). In addition, mathematical models can be used to simulate the outcomes of different management options, allowing farmers and managers to optimize disease control strategies (Gilligan 2008; Cunniffe et al. 2015b; Rimbaud et al. 2018; Fabre et al. 2019). While modeling is not a one-stop solution to the problem of emerging plant pathogens, using mathematical models in conjunction with adaptive management strategies and a robust disease surveillance network can result in more effective disease control.

### Conclusions and Future Directions

Using a spatially-explicit model, we found that the spread of coffee rust is primarily affected by clustering of coffee plants (Figure 3). In addition, clustering interacts with deforestation and fragmentation to influence disease spread (Figures 4, 5). Overall, the models predict a general pattern of highly clustered coffee plots and deforestation leading to greater coffee rust prevalence, with dispersed deforestation increasing individual outbreaks’ predictability.

Our results have important implications for local and regional management practices. Locally, the results of this study show that coffee farms should focus on decreasing the density of coffee plants per coffee plot to prevent coffee rust prevalence, reinforcing the common rust management practice of using shade crops between coffee lines. Regionally, reforestation projects or landscape management decisions should consider that landscapes with higher forest cover tend to have less prevalence of coffee rust and be more resilient against outbreaks. Also, land-use decision makers should consider that the exact degree and location of deforestation matters for disease outbreaks. Higher dispersion of deforestation seems to lead to less variability in rust prevalence, increasing its predictability. Future work should examine how these results play out in real landscapes of coffee plants and forest habitat. In addition, incorporating more biological mechanisms (e.g., wind) into our spatially-explicit model may yield additional insights.

## Code and Data Availability

All code and model outputs are available at https://github.com/Beasley015/QuestCoffeeRustLandscape.

## Acknowledgements

We thank Nick Gotelli and members of the QuEST community at the University of Vermont for helpful feedback on our simulation and analysis.

